# Common and phylogenetically widespread coding for peptides by bacterial small RNAs

**DOI:** 10.1101/030619

**Authors:** Robin C. Friedman, Stefan Kalkhof, Olivia Doppelt-Azeroual, Stephan Mueller, Martiná Chovancová, Martin von Bergen, Benno Schwikowski

## Abstract

While eukaryotic noncoding RNAs have recently received intense scrutiny, it is becoming clear that bacterial transcription is at least as pervasive. Bacterial small RNAs and antisense RNAs (sRNAs) are often assumed to be noncoding, due to their lack of long open reading frames (ORFs). However, there are numerous examples of sRNAs encoding for small proteins, whether or not they also have a regulatory role at the RNA level. Here, we apply flexible machine learning techniques based on sequence features and comparative genomics to quantify the prevalence of sRNA ORFs under natural selection to maintain protein-coding function in phylogenetically diverse bacteria. A majority of annotated sRNAs have at least one ORF between 10 and 50 amino acids long, and we conservatively predict that 188±25.5 unannotated sRNA ORFs are under selection to maintain coding, an average of 13 per species considered here. This implies that overall at least 7.5 ±0.3% of sRNAs have a coding ORF, and in some species at least 20% do. 84 ± 9.8 of these novel coding ORFs have some antisense overlap to annotated ORFs. As experimental validation, many of our predictions are translated according to ribosome profiling data and are identified via mass spectrometry shotgun proteomics. *B. subtilis* sRNAs with coding ORFs are enriched for high expression in biofilms and confluent growth, and two *S. pneumoniae* sRNAs with coding ORFs are involved in virulence. sRNA coding ORFs are enriched for transmembrane domains and many are novel components of type I toxin/antitoxin systems. Our predictions for sRNA coding ORFs, including novel type I toxins, are freely available in a user-friendly format at http://disco-bac.web.pasteur.fr.

## 1 Introduction

Recent technological advances such as tiling microarrays and deep RNA sequencing have led to a new appreciation of bacterial transcription, identifying thousands of new bacterial small RNAs (sRNAs) [50, 19, 23, 42]. Single strains can contain hundreds of sRNAs, including both independent transcripts and extensive transcription antisense to annotated open reading frames (ORFs), even in bacteria lacking the RNA-binding protein Hfq [36, 33, 19]. In virtually all cases, sRNAs contain no annotated ORFs and it is therefore assumed that their primary function is to act as antisense RNAs modulating the expression of other genes [39, 42].

However, gene annotations for short ORFs (usually defined as shorter than 100 or 50 amino acids) are notoriously incomplete and thousands of protein-coding genes remain unannotated in bacteria [46], so many sRNAs could in theory code for functional small proteins. There are several examples of dual-function sRNAs having a regulatory role that also code for an experimentally-validated functional small protein: *E. coli* SgrS encodes the protein SgrT [43], *S. aureus* RNAIII encodes *δ*-hemolysin [52], *B. subtilis* SR1 encodes SR1P [13], and *P. aeruginosa* PhrS encodes an unnamed protein [38]. However, because most known sRNAs have no known antisense regulatory function [23], their primary function could be simply coding for functional peptides.

Small proteins play important roles in bacteria, including quorum sensing, transcription, translation, stress response, metabolism, and sporulation [54, 17]. However, they are difficult to identify by computational or experimental methods. The short sequences have less space for evidence of natural selection, resulting in high levels of statistical noise and false postives, making computational discrimination of coding ORFs smaller than about 50 amino acids difficult [46, 35]. Standard proteomics methods usually utilize gel electrophoresis, which biases towards proteins larger than about 30 kDa and precludes detection of very small proteins [12, 15]. Proteolytic cleavage of some small proteins also results in no peptides of a length detectable by mass spectrometers.

Nevertheless, efforts to identify bacterial short coding sequences have had some success. Proteogenomics, the reannotation of genomes using mass-spectrometry-based proteomics, is a powerful tool for identifying protein-coding genes but still suffers from false negatives, especially for small proteins [15, 40, 28]. Most computational methods applied so far have not taken advantage of sRNA annotations and either used comparative genomics information exclusively [46, 48] or were applied only to a single species [18, 16, 35]. No existing method is ideal for determining the overall number of sRNA coding ORFs. Some comparative genomics methods take into account more information than the *D_n_/D_s_* test, but more complexity can make algorithms more brittle. For PhyloCSF [24], a greater number of parameters to fit can be problematic for small bacterial genomes and this method remains untested on prokaryotes. RNAcode [49] handles multiple alignment issues like insertions and deletions intelligently, but because it does not take into account phylogenetic structure it relies on careful selection of orthologous species to yield relevant results, making it difficult to apply on a large scale. Warren et al. [46] used a clever BLAST-based approach to quickly find new genes, but this is less sensitive than D*_n_*/D*_s_*, which is aware of phylogeny and mutations at the DNA level. Other methods are either ad-hoc and difficult to apply to other species [16] and/or do not incorporate both sequence features and comparative genomics [18].

Short proteins can rarely be predicted with nearly 100% confidence because of limited evidence, but most standard gene annotation tools do not provide an estimated false discovery rate (FDR) for marginal predictions, instead choosing ad-hoc cutoffs for amino acid length or coding score. However, even without confident individual predictions, statistically sound conclusions can be made when considering short ORFs in aggregate; for example, the overall number of ORFs under natural selection to maintain protein-coding potential can be estimated.

To identify short coding sequences in diverse species with high fidelity, algorithms must adapt to composition biases such as GC content, the strength and frequency of Shine-Dalgarno sequence motifs, the availability of closely-related genomes, and the structure of the phylogenetic tree relating these species. We set out to reexamine the assumption that most sRNAs are noncoding by applying simple and adaptable computational and statistical methods to a broad range of bacterial species, paying special attention to controlling for several biases in sRNA ORF sequence properties. We developed a computational method to predict coding ORFs called Discovery of sRNA Coding ORFs in Bacteria (DiSCO-Bac). We then validate the translation of predicted coding sRNA ORFs with experimental data from ribosome profiling and mass spectrometry. We also mine experimental data from various sources to show that many of the resulting small proteins are functional, and a surprising number may be encoded antisense to other RNAs, many of which represent toxin-antitoxin systems.

## 2 Material & Methods

### 2.1 Genomic data collection

Bacterial genomes and ORF annotations were downloaded from Genbank, with accession numbers found in Table 1. Bacterial sRNA annotations were downloaded from BSRD version 1.2 [23].

**Table 1:** Species used in this study and associated Genbank accession numbers for their chromosomal genome.

|  | Species | Genbank version |
| --- | --- | --- |
|  | *Bacillus subtilis* subsp. subtilis str. 168 | AL009126.3 |
|  | *Clostridium acetobutylicum* ATCC 824 | AE001437.1 |
|  | *Escherichia coli* str. K12 substr. MG1655 | U00096.2 |
|  | *Helicobacter pylori* 26695 | CP003904.1 |
|  | *Legionella pneumophila* subsp. pneumophila str. Philadelphia | AE017354.1 |
|  | *Listeria monocytogenes* strain EGD | AL591824.1 |
|  | *Salmonella enterica* subsp. enterica serovar Typhimurium SL1344 | FQ312003.1 |
|  | *Staphylococcus aureus* subsp. aureus N315 | NC_002745.2 |
|  | *Staphylococcus aureus* subsp. aureus NCTC 8325 | CP000253.1 |
|  | *Sinorhizobium meliloti* 1021 | AL591688.1 |
|  | *Streptococcus pneumoniae* TIGR4 | AE005672.3 |
|  | *Streptococcus pyogenes* MGAS5005 | CP000017.1 |
|  | *Streptomyces coelicolor* A3(2) | NC_003888.3 |
|  | *Synechocystis* sp. PCC 6803 | BA000022.2 |

### 2.2 Sequence features

sRNA ORFs between 10 and 50 amino acids in length were found assuming ATG as the only start codon. The 50 amino acid maximum length was applied to account for full-length ORFs that would be found by conventional algorithms but were not included in a particular genome’s Genbank annotations. When an ORF extended past the 3′ end of an annotated sRNA, it was extended until the nearest in-frame stop codon, allowing for inaccurate sRNA 3′ end annotations. sRNA ORFs having in-frame overlap in the same sense as an annotated coding ORF were filtered out. When all intergenic ORFs were considered, no overlap was allowed with annotated coding ORFs, regardless of the frame. The number of ORFs expected by chance (Figure 1B) was generated by shuffling the sRNA sequences one thousand times and counting the resulting ORFs using the same procedure as above. In many instances, multiple start codons could be matched to the same stop codon, corresponding to overlapping in-frame protein sequences. In all predictions of coding ORFs (Figures 3, 4, and 6, these in-frame overlaps were merged, resulting in only the longest ORF, which avoids double-counting coding ORFs. The ORF with the highest coding score was retained in these cases.

**Figure 1:**
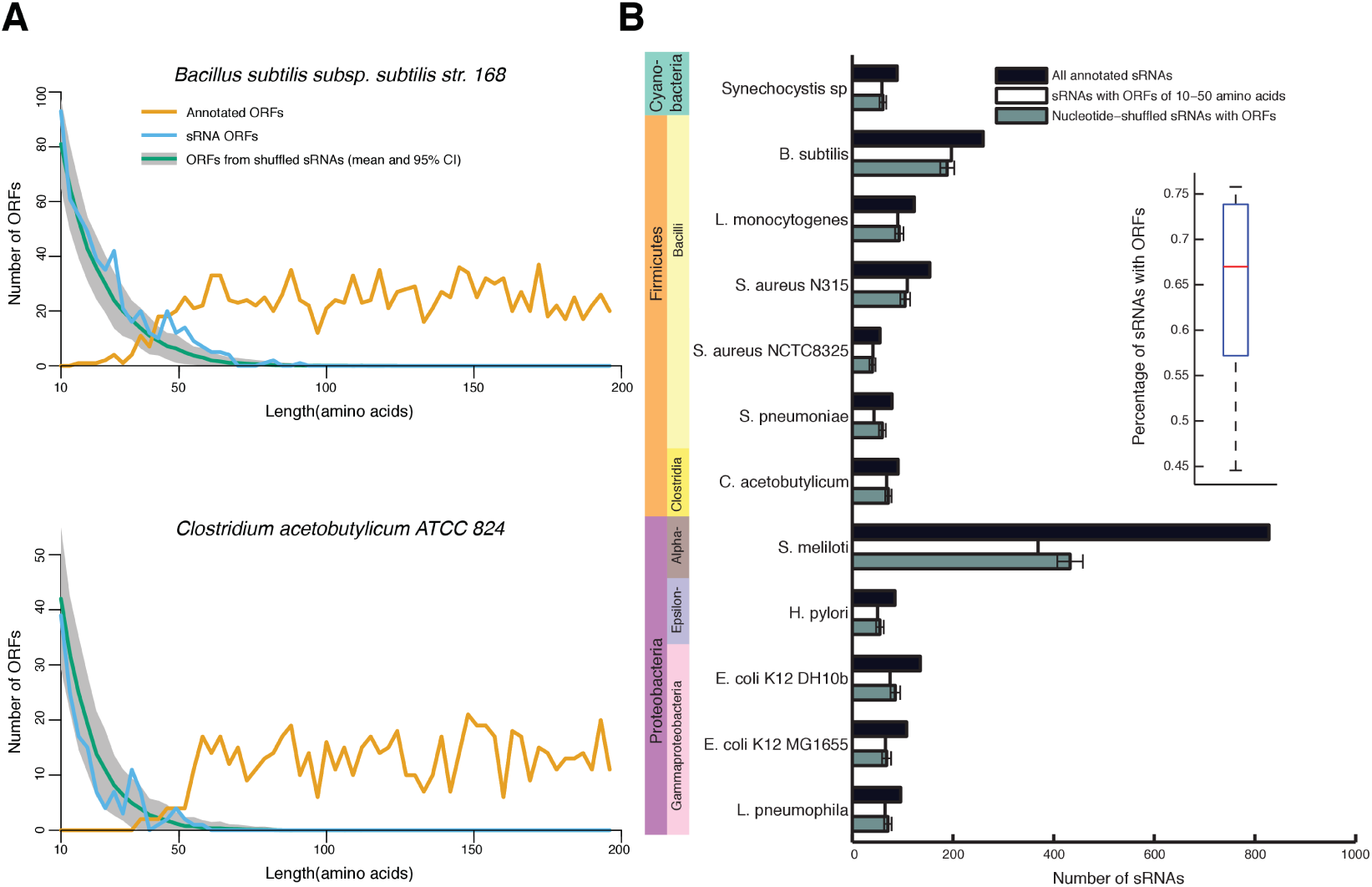
Most sRNAs have at least one potential protein-coding ORF. **(A)** Top: Length distribution of *B. subtilis* 168 ORFs annotated in Genbank (orange) compared to those between 10 and 50 amino acids of length in annotated sRNAs (blue) or those arising by chance in shuffled sRNAs (green). 95% confidence limits based on one thousand shuffles of sRNA sequence are shown in grey. Bottom: The same for *Clostridium acetobutylicum* ATCC 824 ORFs, which have a sharper dropoff around 50 amino acids. **(B)** Number of annotated sRNAs and sRNAs with at least one ORF for 14 species, representing 4 phyla. Phyla are represented as colors on the left, and colors on the right indicate when multiple taxonomic classes are represented within one phylum. As in (A), shuffled sRNA sequences generate a number of ORFs comparable in length to the observed amount. Inset: For each species, the percentage of sRNAs having at least one ORF of between 10 and 50 amino acids in length. Box represents median and first and third quartiles, and whiskers extend to the most extreme values. Most species have ORFs in more than 50% of sRNAs.

Shine-Dalgarno sequence strength (the SD score) was calculated as in Ma & Karlin [25] using their published anti-SD sequences or when unavailable, the corresponding section of the species’ 16S rRNA sequence downloaded from Genbank. For each ORF or mock ORF, ensemble free energies of SD binding were calculated by hybridizing the twenty basepairs upstream of the start codon to the anti-SD sequence using the “hybrid” program from the UNAFold package [27]. Energies reported are at 37 degrees Celsius and in units of -kJ/mol.

Composition bias was calculated using the 69-parameter Z-curve transformation, i.e. phase-specific mononucleotide frequency at each position, and di- and tri-nucleotide frequency at frame zero only [11]. The Z-curve transformation is a numerical representation of the codon bias of an ORF, but also of amino acid frequencies and nucleotide biases such as GC content and purine bias. Z-curve transformations of ORFs and matching intergenic controls were then used to train a logistic regression classifier in Weka version 3-7-9 [14] with a ridge parameter of 10^−8^ yielding scores ranging between zero and one, representing the estimated probability that a sequence is protein-coding. We found that this classifier was more robust and more interpretable than the original Fisher discriminant method used by Gao and Zhang [11]. We skipped any ATG codons when scoring any sequence, because they are, by definition, almost never found in the intergenic mock ORFs.

Density plots in Figure 2 and Supplemental Figure 1 and 3 were smoothed using a gaussian kernel using the *density* function in R with *adjust=0.7.* 95% confidence intervals were ±1.96 times the standard deviation of the control sets.

**Figure 2:**
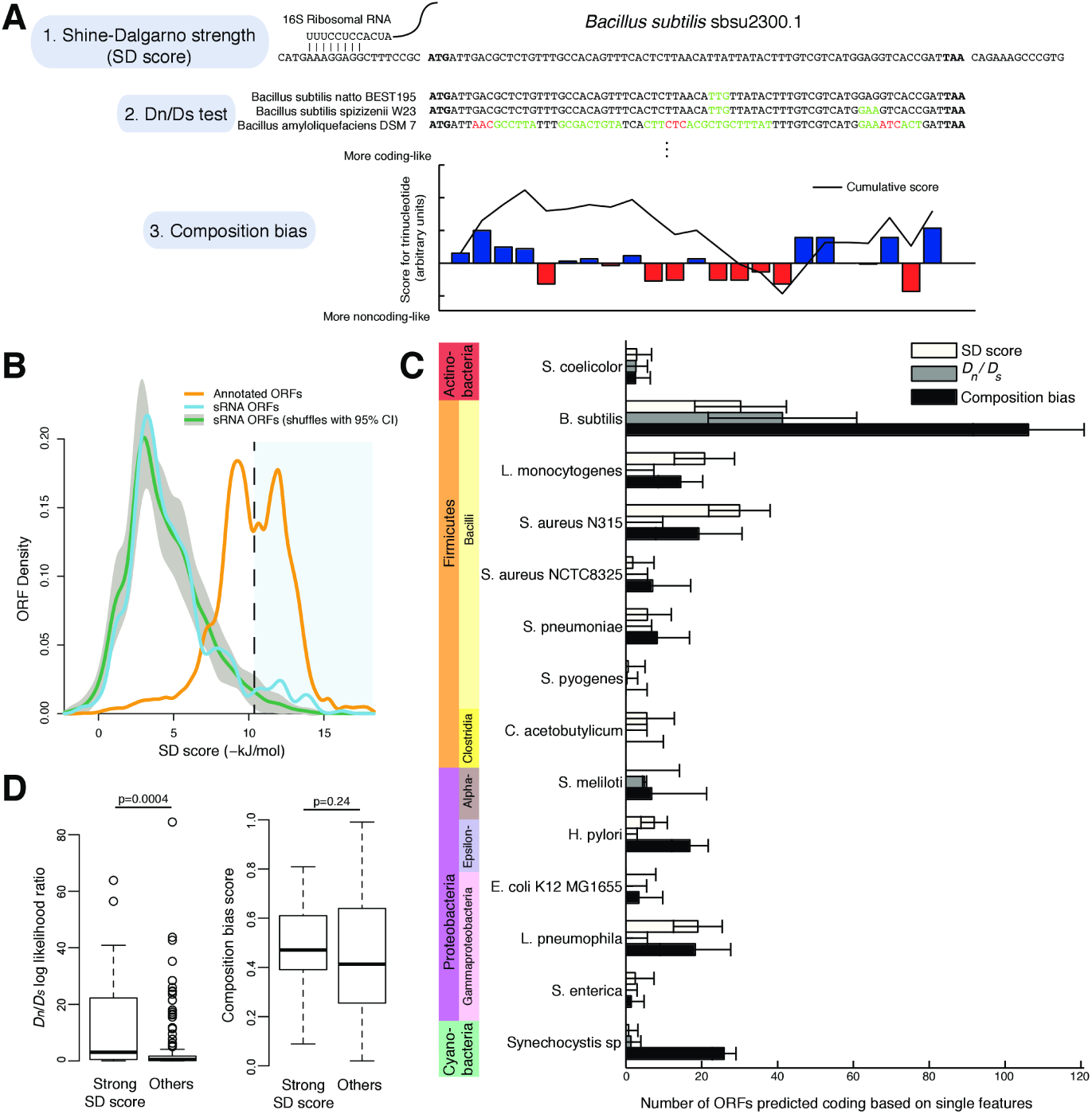
sRNA ORFs have features characteristic of coding ORFs. **(A)** The three features useful for separating coding from non-coding sequences, as illustrated using a *B. subtilis* sRNA that was annotated by tiling array: (1) The Shine-Dalgarno free energy (SD score) measures the free energy of pairing to the 16S ribosomal RNA sequence, which enhances translation to a variable degree depending on the species. (2) The *D_n_/D_s_* test measures whether there is significant conservation on the amino acid level relative to the DNA level by measuring the rate of nonsynonymous and synonymous mutations. For three selected orthologs, synonymous mutations are highlighted in green, nonsynonymous mutations are in red, and start and stop codons are in bold. (3) The composition bias measures phase-specific nucleotide, dinucleotide, and trinucleotide occurrences, learning the difference between coding and noncoding sequence using a logistic regression on training data. The contribution of each codon to the logistic regression score is plotted with bars, with the cumulative score as a black line. **(B)** An example of using a feature to predict coding ORFs in B. *subtilis.* Annotated coding ORFs (orange) have Shine-Dalgarno free energies greater than that expected by chance (green, with 95% confidence limits in gray). Actual sRNA ORFs follow the same distribution, except for an excess of ORFs with free energies more than about -10 kJ/mol (light blue region). **(C)** The number of ORFs predicted as coding based individual features. For each feature, the cutoff with the maximum difference between sRNA ORFs and the background expectation was selected. After correcting empirically for this degree of freedom in selecting the cutoff (see Methods), the number of ORFs predicted coding above background is plotted with 95% confidence intervals. **(D)** Different features sometimes implicate the same ORFs as coding. B. *subtilis* sRNA ORFs were separated into those with SD score stronger than -11 kJ/mol and those with weaker SD score (others). Those with strong SD score (n=27) also had higher Dn/Ds log likelihoods (left panel) but not higher composition bias scores (right panel).

### 2.3 Comparative genomics

Orthologous sequences were selected with a blastn search against all complete bacterial genomes in Genbank with an E-value cutoff of 0.05 and parameters -word_size 7 -gapopen 5 -gapextend 2 -penalty -3 -reward 2. After running blastn for all full-length annotated ORFs in the reference species, species with at least 50% of the maximum number of hits were included as orthologous species; in other words, 50% of the maximum was our ad-hoc threshold for the number of total genes shared with species to be included in the tree. For this analysis, plasmids were included in the search and counted towards the number of hits for a species. To construct a phylogenetic tree, annotated ORFs in the reference sequence were truncated to a maximum of 5000 nucleotides, reblasted against the orthologous species, and aligned using Clustal Omega [37] with default parameters. The alignments were then concatenated and analyzed with the dnaml program of the PHYLIP package using default options [6]. To limit memory usage and running time, we randomly subsampled the sites to yield a maximum of 5 million total nucleotides including all species. For species having more than fifty members of the tree at this stage, we pruned the tree using a greedy algorithm to remove redundant leaves and decrease computational complexity, i.e. we iteratively removed one random species from the pair with the shortest intervening branch length until only fifty leaves remained. At this point, multiple alignments were recreated as above and the phylogenetic tree recalculated using the fifty remaining orthologous species. Any gap positions in the reference organism sequence were spliced out of the multiple alignment, and in-frame stop codons in non-reference species were replaced by gaps. These steps were intended to maximize the amount of data to the *D_n_*/*D_s_* tests while remaining robust to indels and substitutions in the genome assembly and alignment errors. Misaligned sequences statistically add equally to *D_n_* and *D_s_*, thus providing evidence against protein-coding potential, and therefore making the method more conservative with respect to predicting functional coding sequences. *D_n_/D_s_* tests were performed using the codeml program of the PAML package using equal amino acid distance and one site type [53]. Codeml was run once with a fixed omega of one and once with a variable omega, and the reported log likelihood is the difference between the log likelihood of the two models (or zero if the fixed-omega model was more likely than the variable-omega model). We did not perform the *D_n_/D_s_* test on sRNA ORFs overlapping with annotated coding ORFs on either strand, as, in these cases, any signal for selection could not be solely attributed to the sRNA ORF.

To search for homologs to amino acid sequences, command-line BLASTP was used to search the non-redundant protein database (Nr) with the parameters -task blastp-short -evalue 0.25. The search was performed both with and without composition-based statistics (-comp_based_stats 0), and all matches were filtered to be within a two-fold difference in length to the predicted short protein. Any matches with E-value j0.25 in either search were considered homologs. Homologs with descriptions containing “hypothetical”, “uncharacterized”, “unknown”, “unnamed”, “predicted”, “undefined function”, or beginning with “conserved domain protein” were considered to be hypothetical proteins.

### 2.4 Machine learning

All machine learning was performed with Weka version 3-7-9 [14]. When positive and negative sets were not perfectly matched for size, a cost-sensitive classifier was used to ensure an equal weighting of positive and negative training data. For each sequence feature, a unique set of negative controls (which we refer to as “mock ORFs”) was selected to best match relevant properties, and positive controls were sampled from annotated ORFs. These controls were used for statistical tests, plotting figures, and as training data for the machine learning step of coding prediction.

sRNA ORFs having any overlap with an annotated coding ORF were treated separately and were given their own mock ORF sets as described below. The conservation analysis was not performed for these ORFs and was treated as missing data, because the conservation of the annotated ORF would typically have overwhelmed signal for conservation out-of-frame or in the antisense orientation.

For Shine-Dalgarno sequences, the scores of the 20 nucleotides upstream of all annotated ORFs were used as positive controls, as in Ma & Karlin [25]. These sequences were shuffled (i.e. randomly reordered) twenty times and rescored to generate negative control sets controlled for nucleotide composition in each upstream region. The same was done for sRNA ORFs.

For conservation analysis, positive and negative control sets were matched for ORF length and were not shuffled, therefore preserving frame-specific composition and spatial clustering of conservation. Instead, each positive control set consisted of in-frame subsets of annotated ORFs matching the length distribution of sRNA ORFs. Each negative control set consisted of mock ORFs with the same length distribution as sRNA ORFs but randomly taken from “intergenic” regions between annotated coding ORFs, also excluding all sRNA ORFs between 10 and 50 amino acids long. Mock ORFs did not start with an ATG codon, but were constrained not to contain an in-frame stop codon. Twenty sets of positive and negative controls were generated, with each set equal in size and length distribution to the sRNA ORFs. The resulting background estimate of noncoding background is conservative, because the mock ORFs may contain coding ORFs not starting with ATG codons.

Positive and negative controls for the composition bias analysis were the same as those used for the conservation analysis except in the case of sRNA ORFs having overlap with annotated coding ORFs. For each of these sRNA ORFs, twenty mock ORFs were randomly selected with the same overlap length with an annotated ORF in the same orientation (and by implication the same reading frame). For example, an sRNA ORF of 50 nucleotides with its last 10 nucleotides overlapping the 3′ end of an annotated coding ORF would have a negative control set of 20 corresponding mock ORFs having 40 nucleotides in intergenic regions and the last 10 3′ nucleotides overlapping the 3′ end of a randomly selected annotated coding ORF. These mock ORF sets are also required to not contain sRNA ORFs between 10 and 50 amino acids long or in-frame stop codons.

To generate the coding score, we used a two-step classification procedure. First, a logistic regression classifier was trained for composition bias as described above, yielding the composition bias score for each ORF. Next, the SD score, log likelihood of the *D_n_/D_s_* test, and the composition bias score were combined into a training set. As a starting point, the 20 control sets from the conservation and nucleotide composition analysis were used. The positive control sets were matched with the actual SD score of the annotated ORF containing the ORF subset, while the negative control sets were randomly matched to one of the shuffled SD scores. These data were used to train a bagged decision tree classifier using the Weka Bagging class with REPTree base classifiers with default options, but with 100 iterations instead of the default 10. This classifier was used to generate the final coding score of each sRNA ORF, mock ORF (for the calculation of the background distribution), and intergenic ORF. AUC values and ROC curves on training data are for 10-fold cross-validated evaluation. When confidence limits are stated, they represent ± 1.96 times the standard deviation of the negative control sets, i.e., a 95% confidence interval.

Plots in Figure 3A were generated using Weka’s BoundaryVisualizer class using the training data with parameters *r* = 2 and *k* = 5.

**Figure 3:**
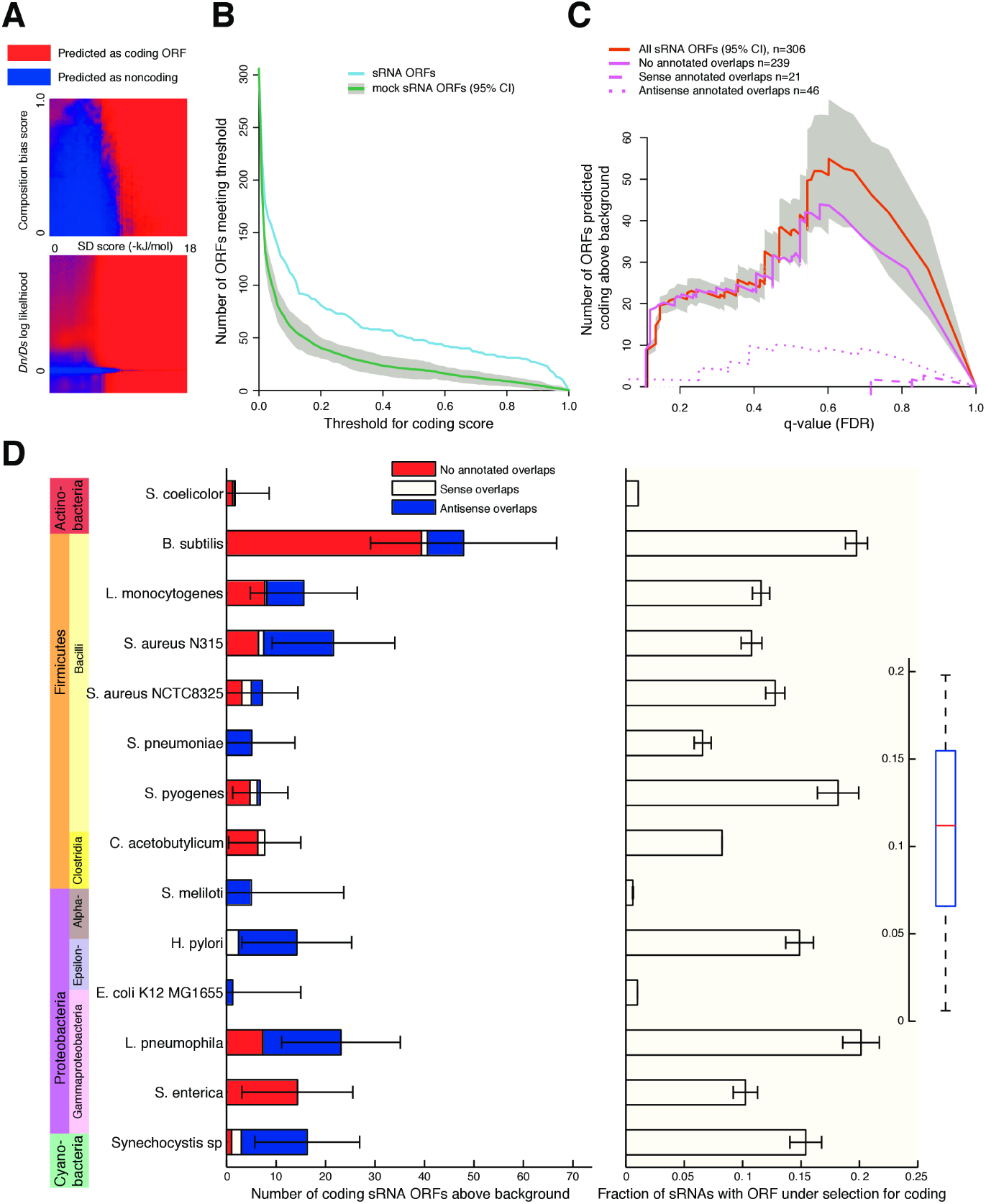
(A) Visualization of a machine-learning classifier for combining features into a single predictive score. A bagged decision tree classifier was trained on *B. subtilis* ORF subsets and mock ORFs, and its output is plotted for each value of SD score and composition bias score (above) or *D_n_/D_s_* (below). For each position, the hidden third feature is subsampled and the classifier output is averaged over these possibilities. **(B)** Number of sRNA ORFs and mock ORFs classified as coding as a function of the coding score threshold in B. *subtilis.* Gray band represents 95% confidence intervals based on 20 mock ORF sets. **(C)** Number of ORFs predicted as coding above background expectation for B. *subtilis.* For each coding score threshold, a false-discovery rate q-value is calculated using the ratio between the sRNA ORFs and mock ORFs plotted in (B). The difference between these two, i.e. the number of ORFs predicted coding above background, is plotted on the y-axis in red, with 95% confidence intervals plotted in gray. The cutoff with the highest sensitivity is marked (dashed black line). The calculation is also made separately for subsets of ORFs having no overlaps with annotated coding ORFs, or those with sense or antisense overlaps. **(D)** Left: Estimated number of ORFs under selection for coding for all species. The numbers predicted for each species were calculated as illustrated in (C) at the most sensitive cutoff and a correction was applied for random fluctuations. Error bars represent 95% confidence intervals on the total estimate, and the breakdown by overlap with annotated coding ORFs is represented by colors. Right: The predicted coding ORFs are sampled to estimate the fraction of sRNAs having at least one coding ORF, and error bars represent 95% confidence intervals. Inset: The fraction of sRNAs having a predicted coding ORF for each species in boxplot form. Box represents first and third quartiles and median; whiskers extend to most extreme values.

### 2.5 Number of sRNA ORFs coding above background

The coding score represents an estimate of the probability that a given ORF is protein-coding, given the assumptions that the training sets accurately represent the parameter distributions for ORFs and noncoding sequences, and that there is an equal number of coding and noncoding ORFs being tested. The latter assumption is problematic, because when we test all sRNA ORFs, far fewer than half may be coding, meaning the coding score is not an accurate probability estimate.

Here, we used the coding score to calculate an empirical false discovery rate estimate. For each species, all in-frame overlapping sRNA ORFs were merged and assigned the largest coding score of the individual ORFs. Let *S* and *M* be the number of sRNA ORFs and mock sRNA ORFs, and *S_T_* and *M_T_* be an estimate of the number of ORFs having coding score greater than a threshold *T*. With the assumption that the distribution of coding scores for noncoding sRNA ORFs is be the same as for mock ORFs, we calculated the empirical false discovery rate (FDR) as

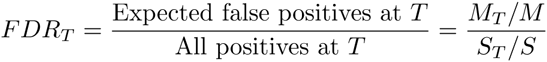
and the *q*-value, the FDR analog of the *p*-value, was by definition

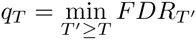

We then estimated the raw number of true coding ORFs, *C*, by

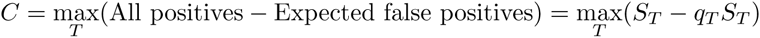

This maximization over all thresholds leaves open the possibility that random fluctuations due to limited sample sizes yield an erroneous signal. This is because any random set of noncoding ORFs will have a transient positive difference *S_T_/S > M_T_/M* at some threshold *T*. Therefore, we made two modifications to this estimate. First we reasoned that the true coding ORFs would be found preferentially at the highest coding scores, so we did not consider any thresholds below the first point at which the coding signal above background was negative, i.e.:

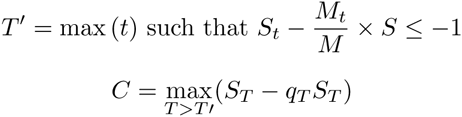

The second modification was to select twenty random subsets of mock ORFs matching the number of true sRNA ORFs to estimate the effect of this random fluctuation, correcting the estimate of coding sRNA ORFs by

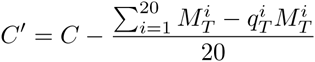
where *M^i^* is subset *i* of the mock ORFs and 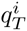 is the calculated as above considering the remainder of the mock ORFs as the false positive distribution. This corrected *C′* is plotted in figure 3D, with the confidence intervals estimated from the standard deviation of the *M^i^* mock ORF subsets.

In species having sRNA ORFs overlapping annotated coding ORFs, the FDR calculation was performed separately for ORFs with no annotated overlap, ORFs with out-of-frame overlap in the sense direction, and ORFs with antisense overlap, yielding three *C′* values. The confidence intervals on sum of the three are based on the standard deviation of the sum of the three *M^i^* mock ORF subsets, one for each category of sRNA ORF.

To estimate the number of sRNAs with at least one coding ORF, we randomly sampled sets of ORFs to be coding 100 times, with each set equal in size to the estimate for the total number of coding sRNA ORFs. This was done separately for ORFs with overlap to annotated coding ORFs in sense and in antisense orientations, and the union of the three sets of coding ORFs were counted.

### 2.6 Intergenic ORFs

When the analysis was expanded to all intergenic ORFs rather than limited to annotated sRNAs, two additional controls were added to eliminate false positives from pseudogenes. First, all ORFs having a *D_n_/D_s_* < 1 with a log likelihood ratio of at least 3, i.e. those having signal for positive selection, were removed as they are likely out-of-frame ORFs in unannotated pseudogenes. Also, all ORFs having a BLAST hit larger than 100 amino acids in length were also eliminated, as they are likely in-frame degenerations of full-length protein.

### 2.7 Ribosome profiling data

Processed ribosome profiling data from [22] was downloaded in processed WIG format from the NCBI Gene Expression Omnibus (Accession GSE35641). Analysis shown is for samples GSM872393 and GSM872397, although similar results were obtained from the other replicates. We counted ORFs as being translated if they had ribosome profiling signal in the correct strand within 3 codons of the start or stop codon, the range reported by Li et al. [22]. ORFs were compared to 20 sets of mock ORFs chosen in the same way as the negative controls for the nucleotide composition analysis. ORFs with out-of-frame overlap to annotated ORFs in the same sense were excluded, because the ribosome profiling data did not have sub-codon resolution.

### 2.8 Proteogenomic analysis of *H. pylori*

Parts of the data and methods were previously published[29], but for convenience we include a complete summary here:

#### 2.8.1 Sample preparation

For validation of predicted novel proteins of *H. pylori* strain 26695 we reanalyzed our in-depth proteome analysis which was especially focused on low molecular weight proteins [Mueller et al., submitted]. Briefly, *H. pylori* strain 26695 was cultured in Ham’s F12 medium (without arginine, Biosera, UK) supplemented with either “light” (12C6, 14N4), “heavy” (13C6, 15N4) or “medium” (13C6, 14N4) isotopically labeled arginine (Cambridge Isotope Laboratories, USA) and 5% (v/v) dialyzed fetal calf serum (FCS) (Thermo Scientific, USA). Cells were harvested and the extracted proteins of the heavy (repG deletion mutant, spiral morphology ([31]), medium (wild type H. pylori strain 26695, spiral morphology) and light arginine (wild type H. pylori strain 26695, coccoid morphology) labeled samples were mixed 1:1:1 (w/w). The protein mixtures were either separated by 1D-SDS-PAGE or, for enrichment of low molecular weight, by gel-eluted liquid fraction entrapment electrophoresis (GELFREE) (5 fractions collecting between 0 and 50 kDa. Proteins were reduced and alkylated and 50% of each sample was digested by endoproteases trypsin and 50% by AspN. Samples were reconstituted with 0.1% (v/v) formic acid for LC-MS/MS analysis.

#### 2.8.2 Nano-uHPLC/nano-ESI analysis

Briefly, proteolytic peptide mixtures were separated on a nano-uHPLC system (nanoAcquity, Waters, Milford, MA, USA) coupled online with an LTQ Orbitrap Velos mass spectrometer (Thermo Fisher Scientific, San Jose, CA, USA). Peptides were trapped and washed for 5 min with 2% acetonitrile containing 0.1% formic acid. Peptide separation was performed using a gradient of 94 min (SDS-PAGE fractions) or 154 min (off-gel fractions) ramping from 2 to 40% acetonitrile, 0.1% formic acid on a C18 column (nanoAcquity UPLC column, C18, 75 *μ*m× 150 mm, 1.7 *μ*m, Waters) with a flow rate of 300 nl/min. The mass spectrometer automatically switched between full scan MS mode (m/z 300-1600, R=60,000) and tandem MS acquisition. Peptide ions exceeding an intensity of 2,000 counts were fragmented within the linear ion trap by collision induced fragmentation. Dynamic precursor exclusion for MS/MS measurements was set to 2 min.

#### 2.8.3 Data Analysis

Proteome Discoverer (version 1.4.1.14, Thermo Scientific., Bremen, Germany) was utilized for peptide identification. The database search engines MS Amanda[4] and Sequest[47] were applied for peptide and protein identification using a concatenated database containing all proteins of H. pylori strain 26695 from NCBI (1596 entries) as well as 89 predicted proteins from an sRNA short ORF with coding *p* > 0.5. The search was conducted allowing a precursor mass tolerance of 10 ppm and a fragment mass tolerance of 0.5 Da. Up to two proteolytic missed cleavages were allowed. Carbamidomethylation of cysteine was defined as fixed modification, whereas oxidation of methionine was set as variable modification for both proteases. AspN specificity was defined to cleave at the N-terminal side of aspartic acid and glutamic acid. A FDR of 5% was applied for peptide and 1% was tolerated for protein identifications. Minimum one unique peptide was required for protein identifications.

#### 2.8.4 Confirmation using synthetic peptides

33 proteolytic peptides of the predicted proteins were synthesized (Thermo Scientific., Bremen, Germany), measured by nano-uHPLC/nano-ESI MS/MS as been descripted above using a 94 min gradient, and were manually compared to spectra obtained by the analyses of the H. pylori samples. Finally, only proteins being identified with a protein FDR below 1% with at least one unique peptide which could be confirmed by a synthetic peptide were reported as novel proteins.

### 2.9 Functional analysis

*B. subtilis* expression data was taken from table S5 of [30]. ORFs in sRNAs or size-matched mock ORFs from intergenic regions were matched to array features if they had any overlap with the annotated feature coordinates. Array features were counted as coding if they contained any sRNA ORF with coding *q* ≤ 0.25. Predicted coding ORFs were then matched to conditions of high expression as annotated by [30] through their array feature. Number of coding ORFs with high expression in a condition expected by chance was calculated based on the mean and standard deviation of 20 sets of mock ORFs or noncoding ORFs equal in size to the predicted coding ORFs. Categories were considered significantly enriched if there was 1.5-fold more predicted coding ORFs than expected based on either control set, with a minimum of 5 ORFs. 95% confidence intervals were calculated using the normal distribution.

Transmembrane domains were predicted by TMHMM [20]. Two negative control sets were used for comparison: first, sRNA ORFs with coding *q* ≤ 0.25 were shuffled 20 times for one control set; second, mock ORFs were created by selecting random fragments of annotated ORFs matching the sRNA ORFs with coding *q* ≤ 0.25 in length.

Type I toxins were taken from BLAST hits in [8]. Toxin amino acid sequences were filtered for only those 10-50 amino acids in length and clustered using CD-HIT v4.6.1 [10], to select representative sequences not more than 90% similar to each other, yielding 114 positive training examples. Negative training examples were sampled from Uniref50 filtered to remove fragments and proteins containing nonstandard amino acids, with length distribution matching the positive control set exactly. 20 negative training examples were selected for each positive example. For each amino acid sequence, physicochemical properties were predicted using various methods: In vitro aggregation, *α*-helix, *β*-strand, and *β*-turn conformation, and helical aggregation propensity were calculated using TANGO version 2.2 [7] with the parameters ct=N nt=N ph=7 te=298 io=0.1. Isoelectric point and hydropathicity were calculated using Bio::Tools::pICalculator and Bio::Tools::SeqStats from Bioperl 1.006923. A random forest classifier was trained on this data using Weka 3-7-9 with parameters -I 10 -K 0 -S 1 -num-slots 1.

Data from [21] were downloaded from the sequence read archive (SRA, accessions SRR064320, SRR064321, and SRR064325). Short RNA reads were trimmed to remove 3′ adapter sequences as in [21], and mapped to the *S. aureus* NCTC 8325 or *B. subtilis* 168 genomes using Bowtie 1.1.0 using the parameters -v 2 - -m 1 –seed 2 –strata –best. Alignments less than 10 bases long were discarded, and alignments less than 20 bases long had at most one allowed mismatch. Coverage of each annotated ORF was calculated using coverageBed from bedtools 2.19.1 [32]. Coverage was normalized to RPKM using the length of each annotated ORF and the total number of mapping reads.

## 3 Results

### 3.1 Most bacterial sRNAs have open reading frames

Hypothesizing that many bacterial sRNAs contain unannotated short ORFs that could code for small proteins, we retrieved sRNA annotations from BSRD [23] and existing ORF annotations from Genbank, focusing first on a gram-positive model species with particularly good sRNA annotations, *Bacillus subtilis* 168. We observed a sharp decrease in annotated coding ORFs shorter than around 50 amino acids (Figure 1A, top). Because many gene finding algorithms use an arbitrary cutoff of 50 amino acids or greater, the handful of shorter annotated ORFs are generally added by hand after their discovery via small-scale experiments. Many less intensely studied bacterial species have an even sharper cutoff around 50 amino acids, as in *Clostridium acetobutylicum* ATCC 824 (1A, bottom). Presumably, this sharp cutoff reflects a technical artifact of automated annotation pipelines rather than an aspect of the true underlying length distribution. For example, reannotation of coding ORFs in yeast based on ribosome profiling recently found translation of 2,869 non-internal ORFs shorter than 50 amino acids [1]. The presence of an artificial cutoff in bacterial ORF lengths suggests that many short coding ORFs remain unannotated. We next asked how many potentially coding ORFs in sRNAs were unannotated, limiting our search to ORFs of between 10 and 50 amino acids in length. Smaller proteins have a lower chance to be functional (for example, only 0.6% of antimicrobial peptides are less than 10 amino acids in length [44]), and longer proteins would likely be identified by modern gene-finding algorithms. We found 506 ORFs in this size range in the 260 annotated *B. subtilis* sRNAs, with 75.8% of sRNAs having at least one ORF (Figure 1A,B). This was roughly the number of ORFs expected by chance based on shuffles of the sRNA sequences (i.e. based on the length and nucleotide composition) (Figure 1A,B). We next examined 14 bacterial species representing 4 phyla and 7 classes, as a diverse representation of species having well-annotated transcriptomes. Each bacterial species had between 22 and 369 sRNAs having at least one ORF, 31%-76% of all sRNAs (Figure 1B). The number of ORFs meeting our size cutoffs in sRNAs was roughly the same for shuffled control sequences in each species, indicating that there has not been significant natural selection against having many sRNA ORFs. Therefore over a broad phylogenetic range, a majority of bacterial sRNAs have at least one ORF with the potential to code for a protein.

### 3.2 Bacterial sRNA ORFs have sequence features predictive of protein-coding function

Most sRNA ORFs are thought not to be translated into protein products, because the presence of an ORF is necessary but not sufficient for expression at the protein level. For example, the Shine-Dalgarno sequence must be accessible to the ribosome for robust translation of most ORFs. Several sequence features have been used to identify ORFs under natural selection to maintain protein coding, separating them from those not likely to be protein-coding [3, 49, 46, 18, 16, 35]. We use three sequence features with wide generality and applicability: The strength of the Shine-Dalgarno sequence (the SD score), conservation at the amino acid level (the *D_n_/D_s_* test), and phase-specific mono- and oligo-nucleotide bias (composition bias) (Figure 2A). Each has varying predictive power depending on the species considered or on the available close orthologs, but is applicable to a wide phylogenetic range of species.

The free energy of pairing between the 16S rRNA and the Shine-Dalgarno sequence (the SD score) is a commonly used predictor of the protein-coding potential of ORFs based on its requirement in some species for strong translation [25]. For some species this feature alone can separate annotated coding ORFs from mock ORFs remarkably well, as in *B. subtilis* (Figure 2B, orange and green lines). sRNA ORFs typically have similar SD scores to randomly shuffled sequences, but for many species a substantial number comprise a tail with stronger SD scores than expected by chance, i.e. an excess of sRNA ORFs that “look like” coding ORFs by SD score (Figure 2B, blue line). This tail of SD scores stronger than expected by chance can be attributed to natural selection acting to preserve translation; therefore, in *B. subtilis*, based only on their SD scores, we can predict that 30 ± 12.5 sRNA ORFs code for proteins (95% confidence interval; Figure 2B, shaded region). The SD score alone was able to predict at least one coding ORF for 10 of the 14 bacterial species tested (Figure 2C, white bars; Supplemental figure 1).

A more direct test for natural selection maintaining protein function is the classic *D_n_/D_s_* log likelihood test, which compares the mutation rate at the amino acid level to that at the nucleotide level. Although there are more sophisticated methods that can perform better in some circumstances [49, 24] we use the classic *D_n_/D_s_* test because it explicitly controls for phylogeny, making it applicable in many contexts, and it is independent of codon bias and nucleotide composition, which can then be explicitly captured in an orthogonal measure. Also, this test is relatively robust to missing data, the choice of orthologous species, and the nature of the selection acting on the sequences. Sequences lost in orthologous species or diverged too far away to align are generally treated as missing data and therefore count neither for nor against an ORF. Therefore it is a conservative test that can be applied in an automated manner to species with diverse phylogenetic tree structures. Applying this test to 14 bacterial species yielded predictions for coding sRNA ORFs above background in 4 species (Figure 2C, grey bars; Supplemental figure 2).

Phase-specific nucleotide bias has been used to successfully predict protein-coding potential in both short and long ORFs [11, 18]. ORFs have biased nucleotide content overall, in specific phases, and other subtle biases such as codon bias. For example, there is a universal bias for purines at the first codon position [2]. Because composition biases differ qualitatively and quantitatively in each species, they must be learned from training data, i.e. real coding ORFs and noncoding sequences. We use a logistic regression method to learn these biases from a subsets of annotated ORFs and mock ORFs in noncoding regions with properties matching the sRNA ORFs in each species (see Methods). The regression then outputs a score for each sRNA ORF representing its likelihood of being coding. Again, sRNA ORFs tend to have higher scores than mock ORFs, with 12 of the 14 bacterial species tested having an excess (Figure 2C, black bars; Supplemental figure 3).

One concern with the nucleotide bias method is that sequences recently acquired via horizontal gene transfer may have different biases and thus may suffer from drastically reduced prediction accuracy. To evaluate the extent of this problem, we examined the SP*β* prophage in *B. subtilis,* a 134 kilobase region having substantially lower GC content than the overall genome (34.6% compared to 43.5%). After training the only on ORF subsets and mock ORFs outside of the SP*β* region, we calculated the nucleotide bias score for 5941 ORF subsets and mock ORFs coming from 234 full-length ORFs within the region, resulting in 81.0% of instances classified correctly (AUC=0.804). By contrast, training on the ORF subsets and mock ORFs in the SP *β* region itself resulted in only a marginal increase to 83.0% of instances classified correctly (AUC=0.877) when using 10-fold cross-validation. This indicates that the nucleotide bias measure should be robust to recent horizontal gene transfer.

The three sequence features used here are complementary because they rely on different information, so it is not surprising that they predict different numbers of coding ORFs for each species. However, presumably many bona fide coding ORFs should score well in multiple features at once, as they are all signs of natural selection for protein-coding potential. Indeed, *B. subtilis* sRNA ORFs with SD scores better than -11 kJ/mol had significantly higher *D_n_/D_s_* log-likelihood scores but not higher composition bias scores compared to sRNA ORFs with worse SD scores (Figure 2D, *p* = 0.0004, 0.24 respectively, two-tailed Mann-Whitney test). This indicates that the three sequence features may be more or less complementary in different combinations and that there is not a trivial way to combine all three into a single score.

### 3.3 Machine learning predicts numerous protein-coding sRNA ORFs in several bacterial phyla

Although the three features may be correlated, they provide complentary evidence, so they must be combined intelligently to maximize their predictive power. Machine learning provides a natural solution for this problem, i.e. training binary classifiers to distinguish between coding ORFs and noncoding sequence. Because each species differs in the phylogeny of available orthologs, history of horizontal gene transfer, codon bias, GC content, and other factors affecting sequence evolution, no single classification scheme can accurately separate coding from non-coding sequence across species. Therefore, we trained individual classifiers on each species, yielding tailored predictions taking into account the predictive power of each sequence feature in each genomic context. We constructed positive training sets based on subsets of annotated coding ORFs, and negative sets based on mock ORFs in likely noncoding sequence, controlling for relevant properties of sRNA ORFs (see Methods). We trained several types of classifiers on these datasets and selected a bootstrap aggregated (bagged) decision tree classifier, which had consistently high performance on all species, for further analysis (Supplemental figure 4). In general, higher scores for each feature increased the likelihood of predicting an ORF to be coding (Figure 3A). However, the classifier takes into account some complexities, such as the fact that an SD sequence is not always required for coding, and that negative log likelihoods for the *D_n_/D_s_* test (i.e. more nonsynonymous than synonymous mutations) can be evidence for coding as well.

When this classifier is applied to sRNA ORFs, it outputs a “coding score” for each ORF between zero and one, which can be interpreted as a probability that an ORF is coding under the assumption that the prior probability of an ORF to be coding is 50%. We estimate the background distribution of coding scores (i.e. scores for noncoding sRNA ORFs) by applying the classifier to the mock ORFs under 10-fold crossvalidation, because they are matched for several relevant properties of the sRNA ORFs. At any selected threshold for coding scores, if more sRNA ORFs meet the cutoff than are expected based on the background distribution, we can attribute the excess to natural selection to maintain coding. Few *B. subtilis* sRNA ORFs meet very stringent thresholds for the coding score, but even fewer mock ORFs do (Figure 3B, right side). As the threshold is relaxed, the number of sRNA ORFs remaining increases faster than the mock ORFs, so more sRNA ORFs can be predicted as coding. However, most ORFs meet the threshold as it appraoches zero and the separation between sRNA ORFs and mocks disappears (Figure 3B, left side). Therefore both the number of sRNA ORFs predicted as coding and the proportion of predictions that are expected to be false positives (the FDR) depend on the coding score threshold. If we want to be more confident in the predictions of individual coding ORFs, we will make fewer overall predictions, while our best estimate of the total number of coding ORFs will be associated with lower confidence in each individual prediction (higher FDR). Figure 3C illustrates this tradeoff for *B. subtilis,* with the estimated false discovery rate (*q* − value) plotted against the number of coding ORFs predicted above background expectation. If we choose the threshold that maximizes sensitivity (i.e. the maximum number of sRNA ORFs predicted as coding, black dashed line in Figure 3C) and apply a correction for the degree of freedom that this adds (see Methods), we predict that there are in total 50 ± 8.2 *B. subtilis* sRNA coding ORFs (95% confidence interval; Figure 3C). When we perform this calculation separately for sRNA ORFs that overlap annotated coding ORFs (either in the sense or antisense direction), we find very little evidence in support of their coding in *B. subtilis* (Figure 3C, broken lines). However, because we do not perform the *D_n_/D_s_* test for these ORFs and because they have other differing properties, we cannot conclude that there are fewer bona fide sRNA coding ORFs that overlap annotated coding ORFs.

Training bagged decision tree classifiers on other bacterial genomes and applying the same test yields between 1 and 23 predicted coding sRNA ORFs, depending on the species. Because the predictive power of the sequence features used varies, we cannot directly compare the number of predictions across species, i.e. we cannot conclude that any species has more bona fide coding sRNA ORFs than another. In addition, some species have more short ORFs previously annotated than others, which are excluded from the analysis. Combining the 14 species considered here, we predict that 188 ± 25.5 (95% confidence intervals) heretofore unannotated small proteins are coded by sRNA ORFs, an average of 13 per species. We believe this to be a very conservative estimate even without taking into account the limited nature of current sRNA annotations, for two main reasons: first, we use only a limited set of sequence features that can at best detect a fraction of cases of natural selection; second, the classifier expects coding ORFs to have similar properties to full-length annotated coding ORFs, while they may be expressed at a lower level on average and conserved in a different manner. For example, the estimate for protein-coding sRNA ORFs based only on the composition bias feature (Figure 2C) is higher for some species than the combined estimate; this is because the SD score and/or *D_n_/D_s_* features hurt the sRNA ORFs in the machine learning classifier more than they help them. Therefore, we can also report a simple, slightly less conservative estimate for the number of coding sRNA ORFs by using only the composition bias feature of 230 ± 28.2, an average of 16 per species.

Because many sRNAs have multiple ORFs, it is not obvious what fraction contain at least one ORF that is under selection to be protein-coding. To estimate this fraction, we used the estimate for the total number of coding sRNA ORFs and randomly chose 100 sets of particular ORFs as coding (Figure 3C, right). For most species, 5-15% of sRNAs had at least one coding ORF, with a median of 11% and a maximum of 20% for *L. pneumophila.* Over all species tested, 7.5 ± 0.3% of sRNAs had at least one coding ORF.

We find almost no evidence in any species for coding of sRNA ORFs overlapping with annotated coding ORFs on the sense strand (Figure 3C, white bars), but there is a substantial number of predictions overlapping coding ORFs on the antisense strand (blue bars). For example, *L. pneumophila,* which has 40 sRNAs with antisense overlap to annotated coding ORFs [51], has 16 ± 5.0 predicted coding sRNA ORFs in this orientation (Figure 3C). Overall, 84 ± 9.8 of the coding ORFs had antisense overlap to annotated ORFs, compared to 91 ± 19 with no overlaps.

### 3.4 Many sRNA ORFs are translated and expressed as peptides

The predictions of protein-coding ORFs reflect evidence for conservation of a protein-coding function, which implies as a pre-requisite expression at the RNA level and protein level. Any annotated sRNA must have evidence for its expression at the RNA level, so all sRNA ORFs have the potential to be translated into peptides. Therefore, as an independent experimental validation of our predictions, we looked for experimental evidence of translation from two types of data: ribosome profiling, showing the translation of transcripts, and mass spectrometry, showing the accumulation of protein to detectable levels.

We used ribosome profiling data for *B. subtilis* 168 and *E. coli* K12 from [22] to annotate translated sRNA ORFs. Looking for signal accumulating on either the start or stop codon of ORFs not overlapping annotated coding ORFs, we found evidence for translation of 132 out of 397 *B. subtilis* sRNA ORFs, compared to on average 62 ± 11 expected by chance (based on the translation of mock ORFs in regions not annotated as coding); in *E. coli* 54 out of 84 ORFs had evidence for translation compared to 42 ± 8.0 by chance (Figure 4A). Because the ribosome profiling data was strand-specific, we could also test the translation of ORFs in the antisense strand to annotated coding ORFs. In this case, the numbers were 5 of 57 ORFs compared to 1.8 ± 3.0 by chance in *B. subtilis,* and 21 of 45 *E. coli* ORFs compared to 3.9 ± 4.2 by chance. In all, this corresponds to 73 and 29 more unannotated sRNA ORFs translated than expected by chance, respectively.

**Figure 4:**
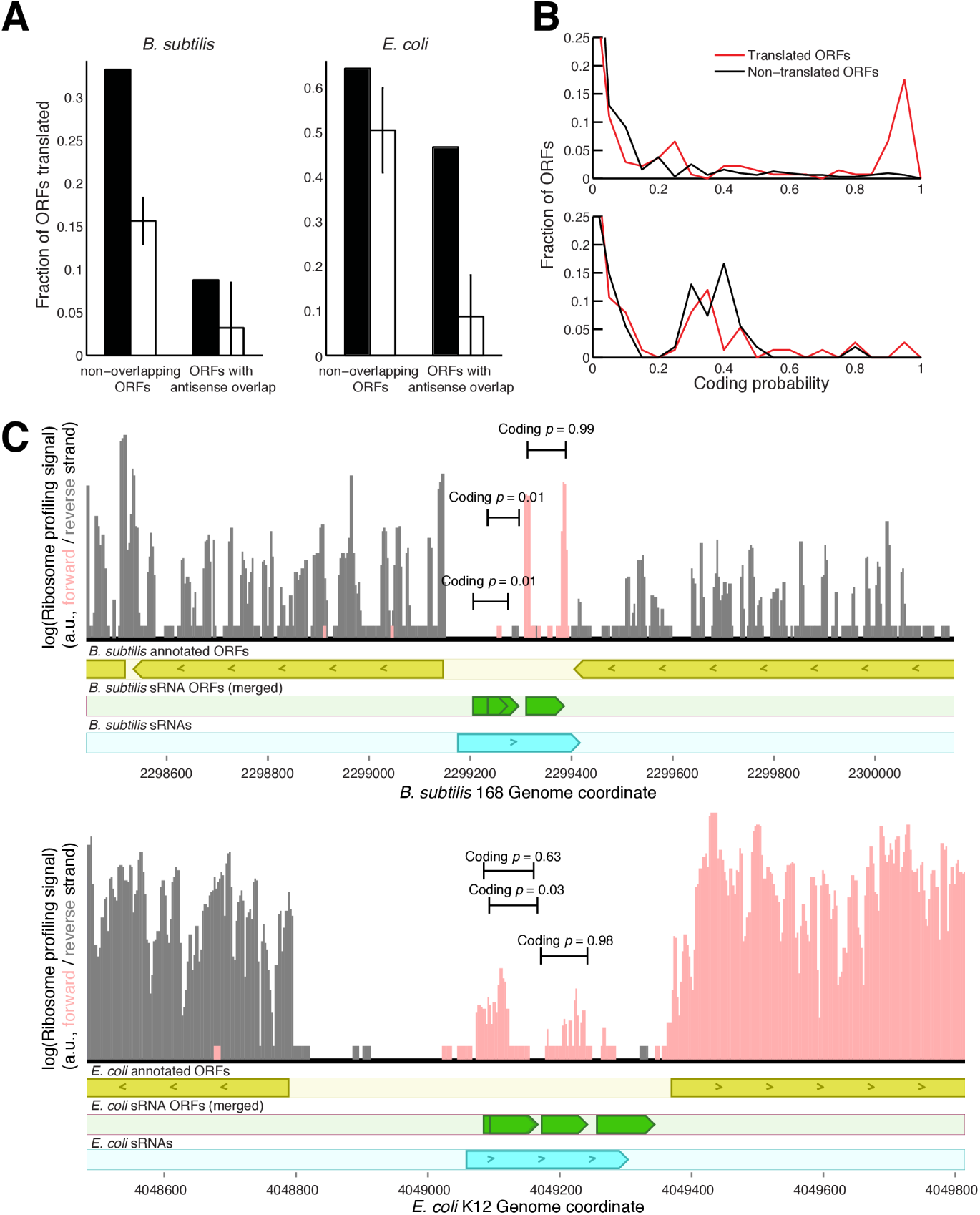
sRNA ORFs with evidence for translation are preferentially predicted to be protein-coding. Ribosome profiling data from [22] was mapped to *B. subtilis* 168 and *E. coli* K12. sRNA ORFs were annotated as translated based only on ribosome profiling signal within three codons of the start or stop codon. **(A)** sRNA ORFs were separated into those independent of annotated ORFs and those antisense to annotated ORFs. The fraction of ORFs translated (black bar) is compared to mock ORFs matched for length and overlap properties (white bars with 95% confidence intervals). **(B)** The predicted coding probability of translated sRNA ORFs is compared to non-translated sRNA ORFs in a histogram for *B. subtilis* (top) and *E. coli* (bottom). **(C)** Top: *B. subtilis* sRNA sbsu2300.1 has three potential coding ORFs, but only one is predicted to be coding. The start and stop codons of this ORF correspond to ribosome profiling peaks, while the ORFs predicted as noncoding do not. Bottom: *E. coli* sRNA seco4050.1 (CsrC) has four ORFs, two of which are predicted as coding and overlap with ribosome profiling peaks. The coding probability can help distinguish between the coding frame for overlapping ORFs, as in the first two in this sRNA.

Some sRNA ORFs with a protein-coding function may not be included in this set because they are only be transcribed and translated under certain condtions, and conversely, translation does not necessarily imply function. Nevertheless, there should be significant overlap between these sets since translation is a prerequisite for protein-encoded function. Indeed, more translated ORFs had more high coding scores than non-translated ORFs, meaning the coding score statistic was able to predict which sRNA ORFs were translated (Figure 4B). This predictive power was not due to the strength of the Shine-Dalgarno sequence alone, implying that selection for protein-coding function reflected in the D_n_/D_s_ test and composition bias was correlated with translation (Supplemental Figure 5). For *E. coli* K12 MG1655, we only predicted 1-2 sRNA ORFs under selection to maintain protein coding, but there was still significant evidence for translation of sRNA ORFs that appeared to be predicted by the coding score. Our methods may be unable to predict these translated protein products with statistical confidence even if they are functional, for example because the SD score has little predictive power in *E. coli* (Supplemental Figure 1), and because translated sRNA ORFs may have different expression profiles and different sequence features compared to annotated ORFs.

Translated ORFs typically had peaks of ribosome profiling signal concentrated at the start or stop codons, as was the case with many full-length annotated ORFs (Figure 4C). Many sRNAs that were previously identified only by high-throughput screens had evidence for translation, such as the *B. subtilis* sRNA sbsu2300.1, which was defined based on tiling array data ([34], Figure 4C, top). Some sRNAs have multiple ORFs between 10 and 50 amino acids long, making the assignment of ribosome profiling coverage to individual ORFs ambiguous. For example, the CsrC noncoding RNA in *E. coli* has ORFs overlapping in different frames (Figure 4C, bottom). In this case, only one ORF under the coverage peak had a coding score of greater than 0.5, showing that the coding score can help to prioritize ORFs for follow-up experiment even when evidence for translation is ambiguous.

### 3.5 Mass spectrometry confirms small proteins from sRNA ORFs

Many sRNA short ORFs could be occasionally translated without accumulation of a protein product to appreciable levels if the protein were quickly degraded. Conversely, detection of sRNA ORF protein products in cells by methods with limited sensitivity would be strong evidence against this possibility. Therefore, we used mass spectrometry specifically designed to find small proteins to search for sRNA ORF products in *Helicobacter pylori.* For the purposes of this search, we included the widest set of sRNA ORF predictions possible, filtering neither for predicted coding score nor potential overlaps with annotated full-length ORFs. This search resulted in 25 peptide hits from 17 sRNA ORFs at an FDR less than 0.05, with 6 hits having an FDR less than 0.01. Despite the statistical significance of these matches, most novel proteins were identified by only a single peptide, making their identification unreliable. The ultimate confirmation of protein identification must be made by matching the observed spectrum to that of a synthetic peptide with the expected sequence. Therefore, we synthesized each of our putative matching peptides. 17 of the synthetic peptides resulted in useable mass spectra, of which 6 tested peptides were validated with a spectrum match. One of the hits, shpy580.1.10, was likely due to an alternative start site of a previously-annotated ORF, i.e. a misannotated or alternative start codon (Supplemental table 1). Another peptide hit in shpy1027.1.1 is for a short sequence shared by an annotated ORF, so may not be considered strong evidence for a novel protein-coding sRNA. However, another peptide comes from an sRNA labeled here as shpy839.1, which is a tmRNA with a known (but unannotated) coding peptide that helps to recycle stalled ribsomes (Figure 5, Supplemental table 1). One other, shpy997.1.2 appears to be a bona fide novel coding peptide arising from an sRNA antisense to an annotated ORF (Figure 5).

**Figure 5:**
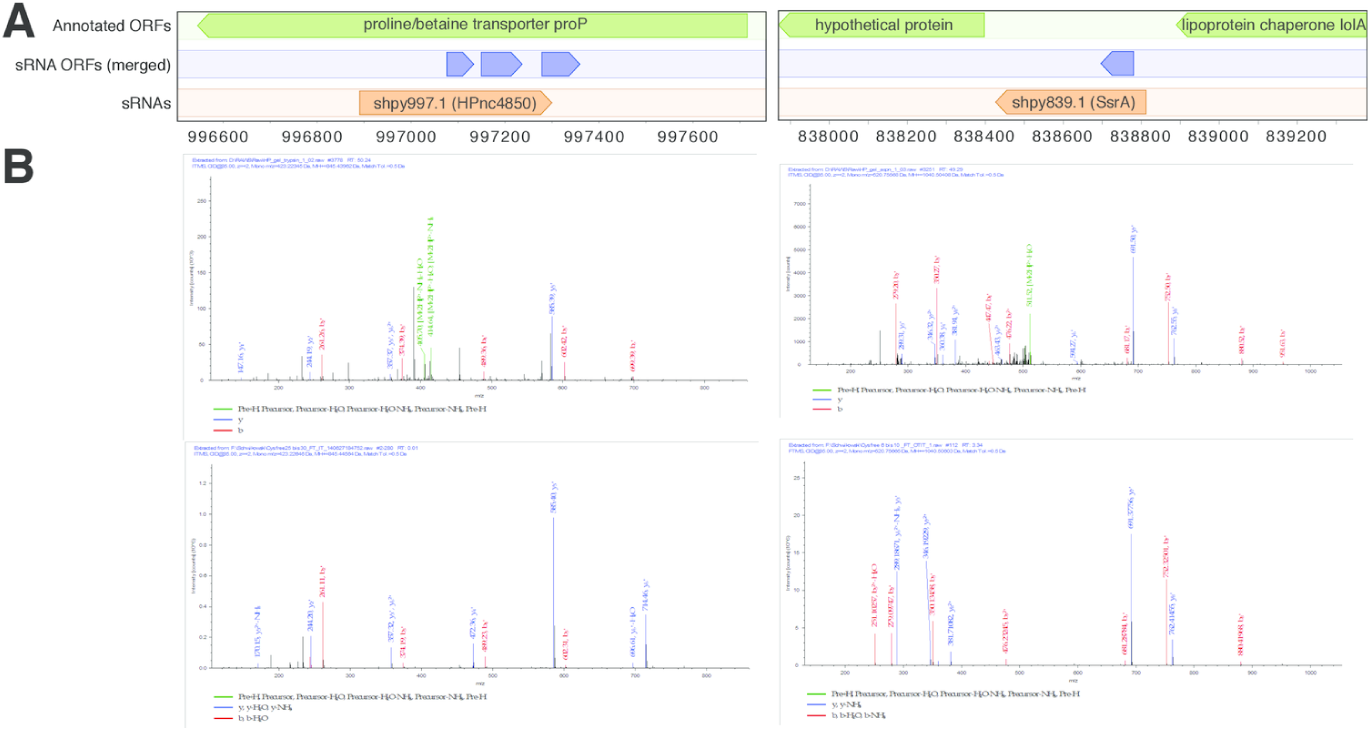
Mass spectrometry confirms sRNA ORFs in *Helicobacter pylori.* A) Two sRNAs are shown in their genomic context, with short ORFs merged for in-frame overlaps and plotted in a separate track. Both sRNAs had a hit in a mass spectrometry search at a false discovery rate of 0.05 (the middle ORF in the case of shpy997.1). B) Validations by synthetic peptide of the peptide/MS matches. On top the spectra from the full shotgun MS/MS experiment are plotted for the hit corresponding to each sRNA ORF. On bottom, the spectra are confirmed with the expected peptide synthesized and run alone on the mass spectrometer.

### 3.6 sRNA ORFs are enriched for certain functions

Some of the sRNAs considered in our study are already known to encode functional peptides in addition to their non-coding functions. For example, the transfer-messenger RNA gene SsrA has tRNA-like properties but also codes for a short peptide, which is rarely annotated. We predicted the coding peptide for SsrA in *S. pneumoniae* at a q-value of 0.10, and in two other species at a coding score *p ≥* 0.5. Most other sRNAs with known coding ORFs were already annotated correctly, for example SgrT in *E. coli* and SR1P/YkzW in *B. subtilis,* so they were skipped by our methods.

To systematically find other annotations for our predicted sRNA coding ORFs, we searched their translated sequences against known proteins using protein BLAST. 21 sRNA ORFs with coding score *p* ≥ 0.5 had a significant match with a similar length and an informative description, including 10 with coding *q ≤* 0.25 (Supplemental Table 2). Many of these BLAST hits were annotated in the same genus as the sRNA ORF, suggesting that they are direct orthologs that have escaped annotation so far. In one case (sbsu22741.1), a *B. subtilis* sRNA ORF matched a type I toxin-antitoxin system annotated in the same species but added to genbank after we performed our analysis. Another notable example is a phenol-soluble modulin that was unannotated in *S. aureus* N315 but is a crucial virulence factor for methicillin-resistant *S. aureus.* Still, the 21 sRNA ORFs with informative BLAST hits leave more than 150 new predicted coding ORFs with no known homolog.

To identify potential functions of our sRNA coding ORFs lacking annotated homologs, we examined a large compendium of *B. subtilis* tiling array data for expression patterns [30]. sRNAs that contained a coding ORF were enriched for high expression in biofilms, stationary phase in minimal media, and other confluent conditions, when compared to sRNAs having ORFs not predicted to be coding, or to mock ORFs (Figure 6).

**Figure 6:**
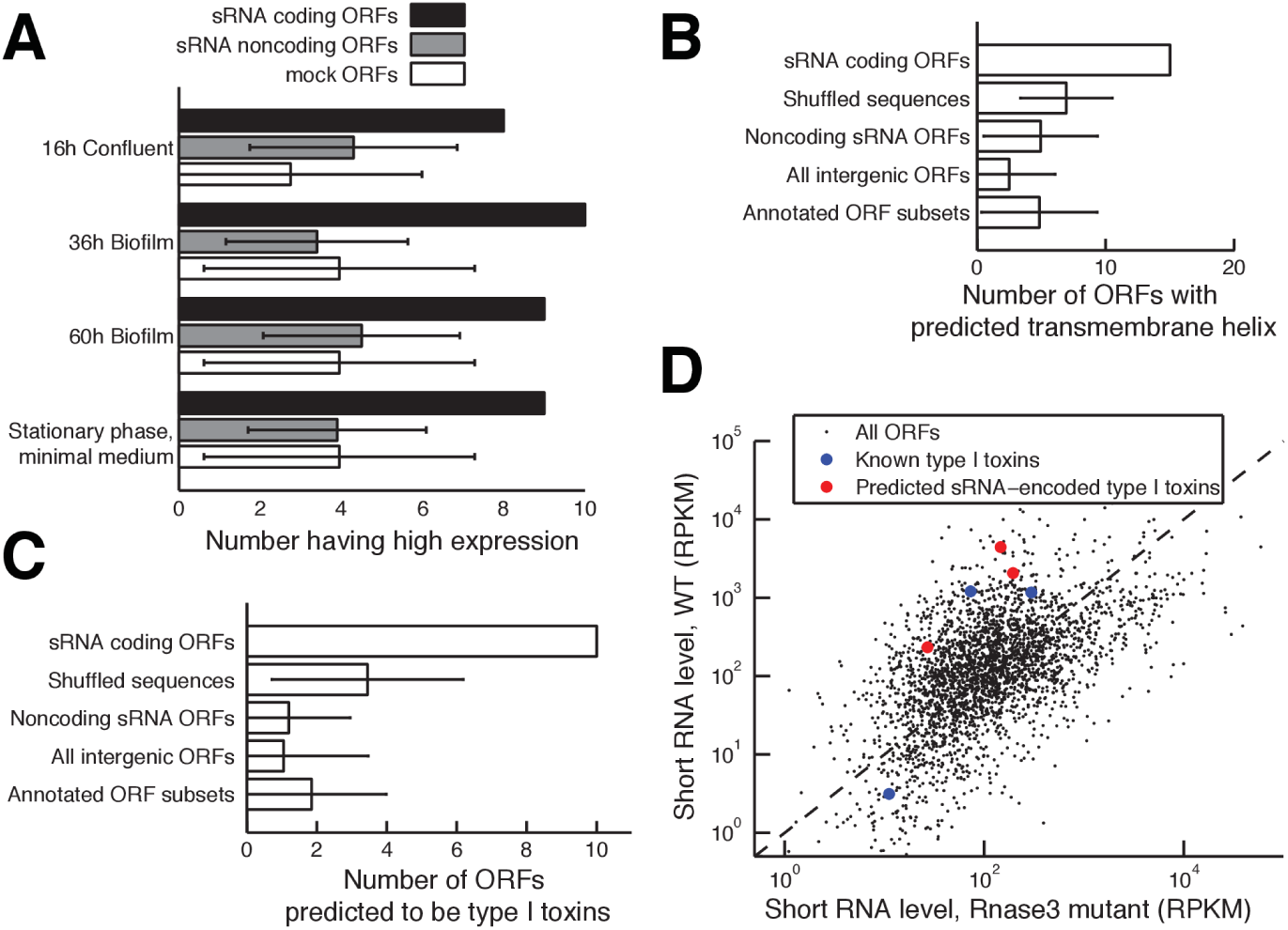
sRNA coding ORFs are enriched for predicted functions including type I toxins. (A) B. subtilis 168 sRNAs containing at least one predicted coding ORF were compared to sRNAs with only noncoding ORFs or mock ORFs. Four growth conditions were enriched for high expression of predicted sRNA coding ORFs (1.5-fold increase in coding ORFs over both controls, minimum 5 sRNA coding ORFs). (B) For all predicted coding ORFs in all species (*q ≤* 0.25, n=120), the number of ORFs with at least one predicted transmembrane helix is compared to the number expected by chance based on their shuffled amino acid sequences, sRNA ORFs with *q* ≥ 0.25, all intergenic ORFs, or size-matched subsets of annotated ORFs. Error bars represent 95% confidence intervals. (C) As in (B), except that the number of ORFs predicted as type I toxins by the physicochemical classifier are plotted. (D) Degradation levels of S. aureus ORFs as measured by sequencing of short RNA fragments are plotted. ORFs subject to degradation of double-stranded RNAs have higher levels in wild-type bacteria (y-axis) than in RNAse III mutant bacteria (x-axis). Known and predicted type I toxins are highlighted (blue and red, respectively).

One of the species we analyzed, *Streptococcus pneumoniae,* was the subject of a recent genome-wide screen for sRNA function in virulence[26]. We searched for coding ORFs in the *S. pneumoniae* sRNAs as defined by Mann et al., resulting in two having coding *q* ≤ 0.25, the R12 and F32 sRNAs. Both had a phenotype when knocked out in the study, with the R12 mutant having reduced fitness in blood and reduced nasopharynx colonization, and the F32 mutant having reduced fitness in Lung infection. The F32 RNA is also known as SsrA, the tmRNA gene with known peptide-encoding function.

### 3.7 Many sRNA ORFs are part of type I toxin/antitoxin systems

An obvious trend in the annotations of the homologs of our predicted coding sRNA ORFs is that 8/20 were membrane-associated, including 3 matches to holins or type I toxin-antitoxin systems, which typically encode membrane-associated small protein toxins (Supplemental Table 2). Therefore we hypothesized that a common function for many sRNA coding ORFs would be to act as membrane-binding proteins and possibly also as toxins. Translated sRNA ORFs with coding q ≤ 0.25 had a predicted transmembrane helix more often than shuffled sequences or equivalent-length regions of annotated ORFs, notably in *B. subtilis, S. aureus* N315, and *S. meliloti* (Figure 6B, Supplemental figure 6). Altogether, the number of coding sRNA ORFs with at least one transmembrane helix in excess of controls was 8.0-13, depending on the type of control, 6.7-11% of the 120 predicted coding ORFs.

To more directly find unannotated toxins, we developed a machine-learning classifier to differentiate type I toxin peptides from non-toxic short proteins based on the physicochemical characteristics of their amino acid sequences, a method adapted from Torrent et al. [41]. We trained a Random Forest classifier on type I toxin peptides found via an exhaustive PSI-BLAST search [8], with length-matched Uniprot sequences as negative controls. This classifier acheived a sensitivity (true positive rate) of 64.3% with a false positive rate of only 3.6% in 10-fold cross-validation of the training data. When applied to predicted sRNA ORFs with coding *q* ≤ 0.25, 10 were predicted type I toxins compared to only 2-4 expected by chance based on shuffled sequences or controls from other ORF types (Figure 6C). Of these 10 predicted toxins, three have annotated holin or type I toxin BLAST hits (sbsu2274.3:2273534-2273824.2, ssau1857.1:1856223-1856978.5, sbsu2679.1:2678645-2679017.7).

If these predicted sRNA coding ORFs are really part of type I toxin-antitoxin systems, their expression should be controlled by the formation of a double-stranded RNA followed by degradation mediated by RNase III. We examined short RNA fragment sequencing data from a recent study for signs of these degradation products in *S. aureus* [21]. Of the three type I toxins reported by Fozo et al. [8], two had very high levels of RNA degradation signal (93rd percentile or above, Figure 6D, Supplemental figure 6). The third was poorly expressed. This signal could not be accounted for by background degradation, as it did not persist in an RNase III mutant (78th percentile or below). We reasoned that novel type I toxins should have similar signal for RNase III dependent degradation. Because we did not predict any confident type I toxins in *S. aureus* sRNAs, we expanded the search to all short ORFs not overlapping annotated ORFs. Three of these ORFs had coding p ≥ 0.5 and were predicted type I toxins. All three had high RNA degradation signal that was dependent on RNAse III to a similar extent (Figure 6D, Supplemental figure 6). Expanding this search to *B. subtilis*, 3/5 predicted type I toxins in sRNA ORFs had high RNA degradation signal (85th percentile and above, supplemental figure 6), although the lack of data from an RNAse III mutant precludes their confident confirmation.

### 3.8 Web Server

We anticipate that this collection of predicted bacterial sRNA coding ORFs with quantified uncertainty will be of broad utility for experimental microbiology. To make these results as accessible as possible, we created a user-friendly web site providing all of our coding ORF predictions for searching or browsing. A parser was written in Python to browse the results of all analyses, formatting and storing them in a couchdb database (http://couchdb.apache.org/). NoSQL technologies were used to dynamically display bioinformatics results. The resulting website is available at the URL http://disco-bac.web.pasteur.fr.

For each species (cf. Table 1), a summary of the coding sRNA ORF search statistics is available. From this first page, the user may view the detailed characteristics including coding p-value for each sRNA ORF, or may expand the search to all intergenic ORFs. In each case, summary figures, for example displaying the ROC curves for prediction accuracy, are also shown. Furthermore, an interactive genome browser [5] was embedded to visualize the genomic context of sRNA ORFs. Tracks show genome position, annotated sRNAs, all sRNA ORFs (with and without in-frame overlaps merged), as well as full-length annotated ORFs. We expect that the accessibility of this data should empower both computationalists and experimentalists to follow up on these results with ease.

## 4 Discussion

We present here a broad survey of protein coding in bacterial sRNAs, to our knowledge, the first of its kind. We combined phylogenetics and known biological effects into a machine learning classifier, a method we call DiSCO-Bac, Discovery of sRNA Coding ORFs in Bacteria. We found that more than half of sRNAs contain canonical ORFs between 10 and 50 amino acids in length, and conservatively at least 7.5% of sRNAs contain ORFs under selection to maintain protein-coding function. In each species considered here, an average of 13 new protein-coding ORFs were predicted in annotated sRNAs. We showed experimental evidence that many of these ORFs were translated (using ribosome profiling data) and that some protein products accumulated to detectable levels (using mass spectrometry). Although few of the predicted protein products had orthologs with annotations, we nonetheless found clues to commonly encoded functions. In *B. subtilis,* sRNAs with coding ORFs were preferentially expressed in biofilms and confluent conditions. Overall, sRNA coding ORFs more often had transmembrane domains than expected by chance. Building a machine learning classifier to find novel type I toxins, we found that sRNA coding ORFs were enriched for toxins, which were associated in *S. aureus* and *B. subtilis* with small RNA degradation products characteristic of control of expression by an RNA antitoxin (Figure 6). Chromosomal type I toxins are often involved in forming bacterial persisters and biofilms [45], consistent with their expression tendencies in *B. subtilis.* We expect that type I toxins constitute a substantial minority of our novel coding ORFs.

Although many are careful to point out that bacterial sRNAs may code for small peptides [50], the term “noncoding” is sometimes used interchangibly, and a trans-acting function for sRNAs is sometimes assumed [42]. Given that more than 50% of sRNAs contain ORFs and up to 60% of *E. coli* sRNA ORFs have some evidence for translation in a single condition (Figure 4A), it is clear that sRNAs should not be assumed noncoding and should not be assumed to have only a trans-acting antisense function. Underscoring this point, for many species, a high fraction of antisense ORFs are predicted to be coding (Supplemental figure 4C). The fact that two sRNAs involved in *S. pneumoniae* virulence, R12 and F32, have confident coding ORFs underscores that the possibility of protein-coding function must always be considered. Of course, there are many examples of sRNAs that have both protein-coding and RNA-level functions, so one does not preclude the other [52, 43, 38, 13]. Antisense transcription is prevalent in many bacteria [21, 42] but these RNAs are even more commonly assumed to function via antisense binding. However, almost half of our newly-identified coding sRNA ORFs had some antisense overlap with annotated ORFs, and these ORFs were found to be commonly translated (Figure 4). For many species, over 20% of antisense ORFs are predicted to be coding (Supplemental figure 4C). This should serve as another reminder that apparent antisense RNA function does not preclude protein-coding potential.

Our survey was enabled by DiSCO-Bac, a flexible machine learning method to find coding sRNA ORFs based on simple sequence features and comparative genomics. Although relatively straightforward, the method is conservative and versatile, making as few assumptions as possible while still being able to incorporate a wide range of evidence. Our use of “mock ORFs” to measure the empirical background distributions of our sequence features is crucial for controlling for several biases that would otherwise strongly skew the analysis, for example, the length distribution and number of ORFs, the GC content of the upstream sequence, higher-order oligonucleotide frequencies, the length of runs of conservation, the effects of overlap with annotated coding ORFs, and the frequency of horizontally transferred sequence. We are careful to guard against double-counting by merging all overlapping ORFs before estimating the number that are coding, to correct for the freedom in fitting the coding score threshold to the data, and to use only a single machine learning algorithm with a single parameter set to guard against researcher degrees of freedom — all of which are common problems in this type of bioinformatic analysis.

Although ultimate proof of protein-coding function for these predicted proteins will await shotgun mass spectrometry targeted towards finding small proteins coupled to genetic and molecular validation, we note that validation at the protein level is extremely difficult. In *H. pylori,* only a single novel short ORF was confidently validated by mass spectrometry using a synthetic peptide, despite dozens of putative hits in the shotgun search. Small proteins are notoriously difficult to detect with shotgun mass spectrometry, because protein purification, electrophoresis, and chromatography all bias towards larger proteins, and because small proteins have few tryptic peptides available to search against. However, we believe more attention paid towards this topic by the proteomics community will yield advances through improved techniques, such as the incorporation of sRNA ORF coding predictions with selected reaction monitoring.

No pre-existing method is ideal for determining the number of sRNA coding ORFs as we have. Some comparative genomics methods take into account more information than the *D_n_/D_s_* test, but more complexity can make algorithms more brittle. For PhyloCSF [24], a greater number of parameters to fit can be problematic for small bacterial genomes and this method remains untested on prokaryotes. RNAcode [49] handles multiple alignment issues like insertions and deletions intelligently, but because it does not take into account phylogenetic structure it relies on careful selection of orthologous species to yield relevant results, making it difficult to apply on a large scale. Warren et al. [46] used a clever BLAST-based approach to quickly find new genes, but this is less sensitive than *D_n_/D_s_*, which is aware of phylogeny and mutations at the DNA level. Other methods are either ad-hoc and difficult to apply to other species [16] and/or do not incorporate both sequence features and comparative genomics [18].

A key advantage of our analysis is the effective aggregation of predictions with weak confidence (high FDR) to make statistically accurate statements about the set of sRNA ORFs as a whole. At first glance, it might seem strange that one can have reasonable confidence in the number of coding ORFs when the confidence in any particular coding ORF prediction is only 50% or less. The situation is analogous to estimating the mean of a probability distribution with a high variance. Each individual observation from the distribution is likely to be far from the mean, but when aggregating over multiple observations, the sample mean converges rapidly to the true expected value as sample size increases, as proven by the law of large numbers and the central limit theorem. There are many instances of biological insight gained by aggregating noisy predictions; for example, the number of human genes that are conserved targets of microRNAs can be quantified despite having low confidence in most individual predictions [9].

One caveat with our method is that it depends on the sRNA annotations, which reduce the amount of ORFs to consider and therefore reduce the background noise and the problem of multiple test correction. For example, *S. aureus* N315 has 154 annotated sRNAs, compared to only 55 in NCTC8325. As a result, we predict 21.6 ± 12.4 *S. aureus* N315 ORFs compared to 7.2 ± 7.2 in *S. aureus* NCTC8325 (Figure 3D). Future additions to sRNA annotations will likely reduce these differences.

We expect that the true number of coding sRNA ORFs is much higher than what we predicted for several reasons: 1) sRNA annotations are based on a subset of published studies which have profiled a subset of potential conditions, and therefore are likely incomplete, 2) the sequence features we used have limited predictive power, for example very few ORFs have strong Shine-Dalgarno sequences in some species, 3) we only consider AUG start codons, while 10.0% of annotated *E. coli* ORFs begin with GUG or UUG, 4) our mock ORFs used for non-coding background calculations likely contain some real ORFs because of missed annotations and nonstandard start codons, 5) we miss unannotated sRNA ORFs smaller than 10 or larger than 50 amino acids, and 6) the machine-learning classifier assumes that sRNA ORFs have similar properties to full-length ORFs, but many bona fide small proteins are likely poorly expressed and/or have different evolutionary histories and thus will have different sequence properties. For example, some sRNA ORFs may have recently evolved *de novo* and therefore would not have orthologs. For all these reasons, we expect that the true number of small proteins encoded by sRNA ORFs is much larger than what we reported, and future improvements to methodology and sRNA annotations will increase these estimates substantially.

It is, however, important to provide predictions now with quantified uncertainty, both to define a lower limit on the size of this class of proteins, and to provide realistic expectations for experimental follow-up. Therefore, we distributed our predictions in a user-friendly web database (http://disco-bac.web.pasteur.fr) combining summary results, individual ORF characteristics, and tools for sorting and visualizing genomic context. These easily-accessible predictions should hasten the experimental characterization of sRNA ORFs. With an average of over a dozen new protein-coding genes predicted in each bacterial species, we expect that future experiments will elucidate novel functions for sRNA ORFs for years to come.

## Acknowledgement

The authors wish to thank Dr. Jeffrey Mellin for inspiration of the project and useful input into the method. RCF was funded by a Pasteur Foundation Postdoctoral Fellowship and ERC grant ERC-2009-AG-232798-HOMEOPITH.

